# Variance-adjusted Mahalanobis (VAM): a fast and accurate method for cell-specific gene set scoring

**DOI:** 10.1101/2020.02.18.954321

**Authors:** H. Robert Frost

## Abstract

Single cell RNA sequencing (scRNA-seq) is a powerful tool for analyzing complex tissues with recent advances enabling the transcriptomic profiling of thousands to tens-of-thousands of individual cells. Although scRNA-seq provides unprecedented insights into the biology of heterogeneous cell populations, analyzing such data on a gene-by-gene basis is challenging due to the large number of tested hypotheses, high level of technical noise and inflated zero counts. One promising approach for addressing these challenges is gene set testing, or pathway analysis. By combining the expression data for all genes in a pathway, gene set testing can mitigate the impacts of sparsity and noise and improve interpretation, replication and statistical power. Unfortunately, statistical and biological differences between single cell and bulk expression measurements make it challenging to use gene set testing methods originally developed for bulk tissue on scRNA-seq data and progress on single cell-specific methods has been limited. To address this challenge, we have developed a new gene set testing method, variance-adjusted Mahalanobis (VAM), that seamlessly integrates with the Seurat framework and is designed to accommodate the technical noise, sparsity and large sample sizes characteristic of scRNA-seq data. The VAM method computes cell-specific pathway scores to transform a cell-by-gene matrix into a cell-by-pathway matrix that can be used for both exploratory data visualization and statistical gene set enrichment analysis. Because the distribution of these scores under the null of uncorrelated technical noise has an accurate gamma approximation, inference can be performed at both the population and single cell levels. As we demonstrate using both simulation studies and real data analyses, the VAM method provides superior classification accuracy at a lower computation cost relative to existing single sample gene set testing approaches.

## 1 Introduction

### 1.1 Single cell transcriptomics

Despite the diversity of cell types and states present in multicellular tissues, high-throughput genome-wide profiling has, until recently, been limited to assays performed on bulk tissue samples. For bulk tissue assays, the measured values reflects the average across a large number of cells and, when significant heterogeneity exists, only approximate the true biological state of the tissue. To address the shortcomings of bulk tissue analysis, researchers have developed a range of techniques for the genome-wide profiling of individual cells [1, 2] with single cell RNA sequencing (scRNA-seq) [3] generating particular scientific interest due to the rapid development of the underlying laboratory techniques, which can now cost-effectively quantify genome-wide transcript abundance for thousands to tens-of-thousands of cells. Single cell genomic assays, in combination with techniques that infer transcription rates [4], spatial information [5] or temporal dynamics [6,7], provide scientists with a detailed picture of cellular biology. Such cell-level genomic resolution is especially important for the study of tissues whose structure and function is defined by complex interactions between multiple distinct cell types that can occupy a range of phenotypic states, e.g., the tumor microenvironment [8, 9], immune cells [10, 11], and the brain [12]. Important scientific questions that can be addressed by single cell transcriptomics include the identification and characterization of the cell types present within a tissue [13, 14], the discovery of novel cell sub-types [15], the analysis of dynamic processes such as differentiation [7], or the cell cycle [16], and the reconstruction of the spatial distribution of cells within a tissue [5].

### 1.2 Single cell analysis challenges

Although single cell data provides unprecedented insights into the structure and function of complex tissues and cell populations, technical and biological limitations make statistical analysis challenging [17]. Single cell methods analyze very small amounts of genomic material, leading to significant amplification bias and inflated zero counts relative to bulk tissue assays [18]. Single cell-specific approaches for quality control, normalization and statistical analysis (e.g., zero-inflated models) only partially address these challenges [19, 20]. In addition to the challenges of increased noise and missing data, important biological differences exist between bulk tissue and single cell data. As the average over a large number of cells, bulk tissue measurements are typically unimodal and, in many cases, approximately normally distributed. In contrast, single cell data sets reflect a heterogeneous mixture of cell types and cell states resulting in multi-modal and non-normal distributions [18]. The diverse mixture of cell types and states found in complex tissues also leads to significant differences in gene expression patterns between bulk tissue and single cell data. As evidenced by projects such as the Human Protein Atlas (HPA) [21], gene activity measured on bulk tissue samples can differ substantially from the activity occurring within the cell subpopulations comprising the tissue. Figure 1 provides a simplified illustration of the marginal and joint distribution characteristics of single cell and bulk tissue gene expression data. In this figure, the marginal distribution is represented by density plots for a single gene while the joint distribution is represented by covariance matrices. Collectively, the distributional differences between single cell and bulk tissue genomic data make it challenging to successfully analyze single cell expression data using methods originally developed for bulk tissue, which assume non-sparse data and lower levels of technical noise.

**Figure 1:**
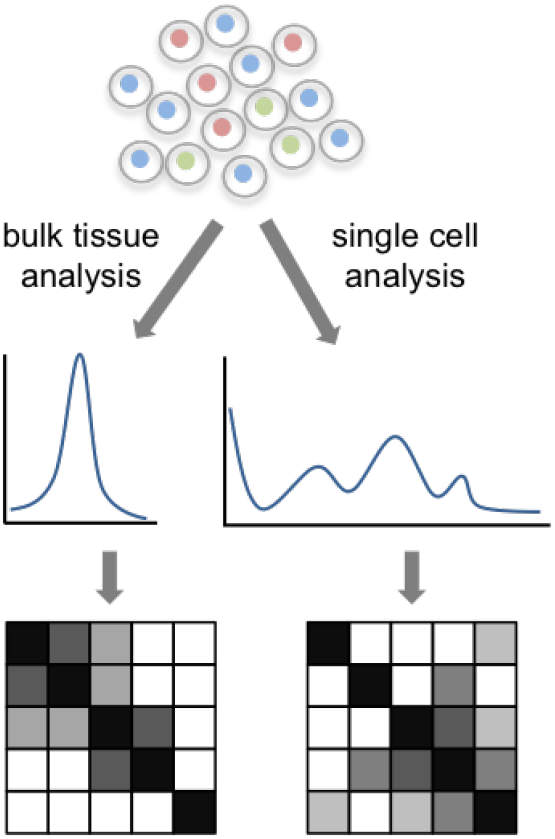
Features of bulk tissue vs. single cell distributions.

### 1.3 Gene set testing of single cell data

Although high-dimensional genomic data provides a molecular-level lens on biological systems, the gain in fidelity obtained by testing thousands of genomic variables comes at the price of impaired interpretation, loss of power due to multiple hypothesis correction and poor reproducibility [22–25]. To help address these challenges for bulk tissue data, researchers developed gene set testing, or pathway analysis, methods [25,26]. Gene set testing is an effective hypothesis aggregation technique that lets researchers step back from the level of individual genomic variables and explore associations for biologically meaningful groups of genes. By focusing the analysis on a small number of functional gene sets, gene set testing can substantially improve power, interpretation and replication relative to an analysis focused on individual genomic variables [22–25]. The benefits that gene set-based hypothesis aggregation offers for the analysis of bulk tissue data are even more pronounced for single cell data given increased technical variance and inflated zero counts.

Gene set testing methods can be categorized according to whether they support supervised or unsupervised analyzes (i.e., test for association with a specific clinical endpoint or test for enrichment in the variance structure of the data), whether they provide results for each sample or for an entire population, whether they test a self-contained or competitive null hypothesis (i.e., the *H*_0_ that none of the genes in the set has an association with the outcome or the *H*_0_ that the genes in the set are not more associated with the outcome than genes not in the set) and whether they test each gene set separately (uniset) or jointly evaluate all sets in a collection (multiset). In this paper, we focus on single sample gene set testing methods, i.e., those that compute a cell-specific statistic for each analyzed gene set to transform a cell-by-gene scRNA-seq matrix into a sample-by-pathway matrix. This class of techniques is of particular interest because the cell-level pathway scores can be leveraged for both exploratory data visualization, e.g., shading of cells in a reduced dimensional plot according to inferred pathway activity, as well as the full range of population-level statistical gene set tests, i.e., supervised or unsupervised tests of either the uniset or multiset flavor.

Existing single sample gene set testing methods can be grouped into three general categories: random walk methods, principal component analysis (PCA)-based methods and z-scoring methods. Random walk methods (e.g., GSVA [27] and ssGSEA [28]) generate sample-level pathway scores using a Kolmogorov-Smirnov (KS) like random walk statistic computed on the gene ranks within each sample, often following some form of gene standardization across the samples. PCA-based methods (e.g., PAGODA [29] and PLAGE [30]) perform a PCA on the expression data for each pathway and use the projection of each sample onto the first PC as a sample-level pathway score. Z-scoring methods (e.g., technique of Lee et al. [31], scSVA [32], and Vision [33]) generate pathway scores based on the standardized mean expression of pathway genes within each sample. While these methods have proven effective for the analysis of bulk expression data, with GSVA and ssGSEA among the most popular techniques, the application of these methods to scRNA-seq data is limited by three main factors: poor classification performance in the presence of sparsity and technical noise, lack of inference support on the single cell level, and high computational cost (esp. for the random walk methods when the number of samples/cells is large).

GSVA, ssGSEA, PLAGE and the Lee et al. z-scoring methods were all developed for the analysis of bulk gene expression data and were therefore optimized for, and evaluated on, non-sparse data with moderate levels of technical noise. Although scSVA and Vision are both targeted at single cell expression data, they are methodologically similar to the Lee et al. z-scoring technique and make no special provision for sparsity or elevated noise. As we demonstrate through simulation studies later in the manuscript, these methods all have poor classification performance relative to the VAM technique on sparse and noisy data, i.e., they are not able to effectively identify cells whose transcriptomic profile is enriched for specific pathways. In contrast to the other existing single sample methods, PAGODA was designed for single cell analysis and specifically addresses the scRNA-seq features of sparsity and technical noise. In the case of PAGODA, however, the primary focus is an unsupervised and population-level analysis; the generation of sample-level scores is a secondary output which lacks inference support and, relative to the random walk and z-scoring approaches, is particularly poor at identifying cells with elevated expression of specific pathways. The practical utility of the PAGODA method is also hindered by a fragile installation procedure (we were unable to install it successfully), the requirement for a specialized normalization process and lack of direct integration with popular scRNA-seq frameworks like Seurat [34, 35].

Although the pathway scores generated by the z-scoring methods should have a standard normal distribution when the expression data follows an uncorrelated multivariate normal distribution, this distributional assumption does not hold for sparse scRNA-seq data. Neither the random walk nor the PCA-based method generate scores with a well characterized null distribution. While the lack of a null distribution does not prevent the cell-specific scores generated by these techniques from being used for visualization or as predictors in regression models, it does preclude cell-level inference and the use of scores as dependent variables in parametric models.

Given experimental and cost constraints, most bulk gene expression data sets have sample sizes in the hundreds; bulk data sets with more than one thousand samples are rare. Single cell data sets, by contrast, typically profile thousands of cells and data sets containing tens-of-thousands to hundreds-of-thousands of cells are becoming increasingly common. These large sample sizes make computational cost an important factor, especially for techniques that are used in an exploratory and interactive context. Relative to the VAM approach, all of the existing single sample methods have significantly worse computational performance on even small (2000 cells, 500 genes) data sets. For very large scRNA-seq data sets (i.e., 100,000+ cells), the use of methods like GSVA and ssGSEA will be impractical for most users.

## 2 Methods

### 2.1 Variance-adjusted Mahalanobis (VAM)

The VAM method generates cell-specific gene set scores from scRNA-seq data using a variation of the classic Mahalanobis multivariate distance measure [36]. VAM takes as input two matrices:

- **X**: *n* × *p* matrix that holds the positive normalized counts for *p* genes in *n* cells as measured via scRNA-seq. As detailed in Section 2.4 below, VAM provides direct support for both Seurat [35] normalization techniques: log-normalization (i.e., log of 1 plus the unnormalized count divided by an appropriate scale factor for the cell) and the SCTransform method [37]. Other scale factor-based normalization techniques that are equivalent to Seurat log-normalization (e.g., normalization supported by the Scater framework [19]) can also be used.
- **A**: *m* × *p* matrix that represents the annotation of *p* genes to *m* gene sets as defined by a collection from a repository like the Molecular Signatures Database (MSigDB) [38] (*a*_*i*,*j*_ = 1 if gene *j* belongs to gene set *i*).

VAM generates as output one matrix:

- **S**: *n* × *m* matrix that holds the cell-specific scores for each of the *m* gene sets defined in **A**.

Given **X** and **A**, VAM computes **S** using the following steps:

1. **Estimate technical variances**: Let 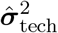 be a length *p* vector holding the technical component of the sample variance of each gene in **X**. For the VAM-Seurat integration, two approaches are supported for computing 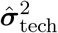 depending on whether log-normalization or SCTransform is employed (see Section 2.4 below for details). Similar variance decomposition approaches are supported by other scRNA-seq normalization pipelines (e.g., Scater [19]). VAM can also be used under the assumption that the observed marginal variance of each gene is entirely technical. In this case, 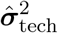 is simply estimated by the sample variances of each gene in **X**.
2. **Compute modified Mahalanobis distances**: Let **M** be an *n* × *m* matrix of squared values of a modified Mahalanobis distance. Each column *k* of **M**, which holds the cell-specific squared distances for gene set *k*, is calculated as:

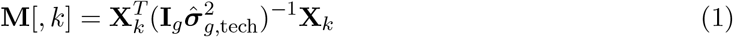

where *g* is the size of gene set *k*, **X**_*k*_ is a *n*×*g* matrix containing the *g* columns of **X** corresponding to the members of set *k*, **I**_*g*_ is a *g* × *g* identity matrix, and 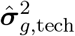 holds the elements of 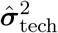 corresponding to the *g* genes in set *k*.
3. **Compute modified Mahalanobis distances on permuted X**: To capture the distribution of the squared modified Mahalanobis distances under the *H*_0_ that the normalized expression values in **X** are uncorrelated with only technical variance, the distances are recomputed on a version of **X** where the row labels of each column are randomly permuted. Let **X**_**p**_ represent the row-permuted version **X** and let **M**_**p**_ be the *n* × *m* matrix that holds the squared modified Mahalanobis distances computed on **X**_**p**_ according to (1).
4. **Fit gamma distribution to each column of M_p_**: A separate gamma distribution is fit using the method of maximum likelihood (as implemented by the *fitdistr()* function in the MASS R package [39]) to the non-zero elements in each column of **M**_**p**_. Let 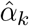 and 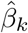, *k* ∈ 1, …, *m* represent the gamma shape and rate parameters estimated for gene set *k* using this procedure. As detailed in Section 2.3, the normal *χ*^2^ approximation for standard squared Mahalanobis distances does not hold for the values generated according to (1), however, the null distribution of these values can be well characterized by a gamma estimated on each column of **M**_**p**_. Note that if computational efficiency is a major concern, the gamma distributions can be fit directly on **M** to avoid the cost of generating **X**_**p**_ and **M**_**p**_; this will impact the power to detect deviations from *H*_0_ but will not inflate the type I error rate.
5. **Use gamma cumulative distribution function (CDF) to compute cell-specific scores**: The cell-specific gene set scores are set to the gamma CDF value for each element of **M**. Specifically, each column *k* of **S**, which holds the cell-specific scores for gene set *k*, is calculated as:

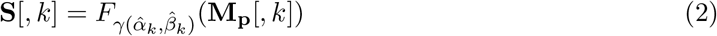

where 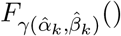 is the CDF for the gamma distribution with shape 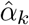 and rate 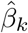. Under the *H*_0_ of uncorrelated technical noise, valid p-values can be generated by subtracting the elements of **S** from 1. Section 2.3 explores the statistical properties of the elements of **M** and inference using p-values generated via **1** − **S** in greater detail.

The use of 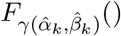 to generate the elements of **S** has several important benefits in addition to support for cell-level inference. First, it transforms the squared modified Mahalanobis distances for gene sets of different sizes into a common scale, which is important if values in **S** are used together in statistical models, e.g., as regression predictors. Second, it generates a statistic that is bound between 0 and 1 and is robust to very large expression values, i.e., the CDF converges quickly to 1 as the squared distances increase. Such robustness is particularly important for the analysis of noisy scRNA-seq data; many existing scRNA-seq analysis methods such as SC-Transform artificially clip normalized data to eliminate extreme values. Lastly, the fact that the distribution of values is often bimodal with most values close to 0 or 1 improves the utility of **S** for both visualization and statistical modeling.

### 2.2 Comparison of VAM and the standard Mahalanobis distance

For the scenario represented by (1), the squared Mahalanobis distance is normally defined as:

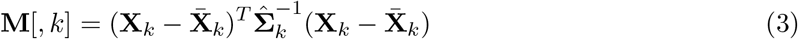

where 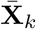 is a matrix whose rows contain the mean values of the columns of **X**_*k*_ and 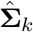 is the estimated sample covariance matrix for **X**_*k*_. There are two important differences between the modified Mahalanobis distance in (1) and the standard Mahalanobis distance in (3):

1. The standard Mahalanobis distance uses the full sample covariance matrix whereas the modified Mahalanobis distance accounts for just the technical variance of each gene and ignores covariances.
2. The standard Mahalanobis measure computes the distances from the multivariate mean whereas the modified Mahalanobis distance in computes distances from the origin.

A key feature of the VAM method, and the basis for the “variance-adjusted” portion of the name, is the use of 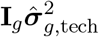 instead of the sample covariance matrix included in the typical formulation of the Mahalanobis distance. The practical impact of this change is that deviations in directions of large estimated technical variance are discounted (i.e., larger deviations are expected due to the higher variance) but deviations in directions of large biological variation (or covariance) are not discounted (i.e., these deviations are not expected if the variation in expression is purely technical).

Use of the origin instead of the multivariate mean in (1) generates a more biologically meaningful distance measure for scRNA-seq data. With the standard Mahalanobis distance, it is possible for samples whose elements are all above the mean, all below the mean or a mixture of above and below to have the exact same distance value. Computing distances from the origin for positive data eliminates this ambiguity: larger distances correspond to larger positive sample values, i.e., elevated gene expression in the cell, and a distance of 0 corresponds to lack of expression in all genes. Measuring distances from the origin will also assign more extreme values to sets whose members show coordinated expression. When distances are measured from the multivariate mean, it is not possible distinguish between sets with a mixture of up and down-regulated genes and sets whose members show coordinated expression. Prioritizing coordinated expression is advantageous since such pathways are usually more biologically interesting. As a simple example, imagine a two gene set with mean (1, 1) and identity covariance matrix. For this set, cells with expression values of (0, 0), (2, 0), (0, 2), and (2, 2) all have the same squared Mahalanobis distance of 2 when distances are measured from the multivariate mean. By contrast, the squared distance from the origin for these cells is 0, 4, 4, and 8, which better reflects the combined expression of these genes. It should be noted that the difference between the mean and origin will be minor for the large number of geens in an scRNA-seq data set that have mean values very close to 0.

### 2.3 Statistical properties of VAM

If the values in **X**_*k*_ follow a multivariate normal distribution, the squared Mahalanobis distances computing according to the standard definition in (3) can be approximated by a *χ*^2^ distribution with *g* degrees-of-freedom, where *g* is the size of gene set *k*. If 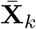 is replaced by the **0** vector in (3), the resulting squared distances are instead approximated by a non-central *χ*^2^ distribution with *g* degrees-of-freedom and non-centrality parameter 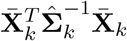.

The modified squared Mahalanobis measure used by VAM and defined in (1) can also be approximated by a non-central *χ*^2^ distribution under the *H*_0_ of uncorrelated technical noise if the data in **X**_*k*_ is not too sparse, i.e.,~ 50% or fewer of the elements are zero, and the non-zero values in **X**_*k*_ have an approximately normal distribution. Figure 2 illustrates the density estimate for values computed using (1) on scRNA-seq data simulated under the *H*_0_ of uncorrelated technical noise for sparsity values of both 0.5 and 0.8 (see Section 2.5 for more details on the simulation model, which assumes a log-normal distribution for the non-zero elements in **X**_*k*_). Figure 2 also includes the density for the non-central *χ*^2^ distribution with the appropriate degrees-of-freedom and non-centrality parameter. As shown in this figure, the non-central *χ*^2^ distribution provides an accurate approximation for a sparsity of 0.5, panel a), but overestimates the mean and significantly underestimates the variance of the squared distances when the sparsity increases to 0.8, panel b). Given the poor fit of a non-central *χ*^2^ distribution for realistic sparsity levels, we instead model the null distribution of elements in **M** by a gamma distribution whose parameters are estimated via maximum likelihood as described in Section 2.1 above. As shown in Figure 2, the estimated gamma distribution provides a very good fit for the observed squared modified Mahalanobis distances at both the 0.5 and 0.8 sparsity levels. The type I error control and power provided by the estimated gamma distribution is detailed in Section 3.1 below.

**Figure 2:**
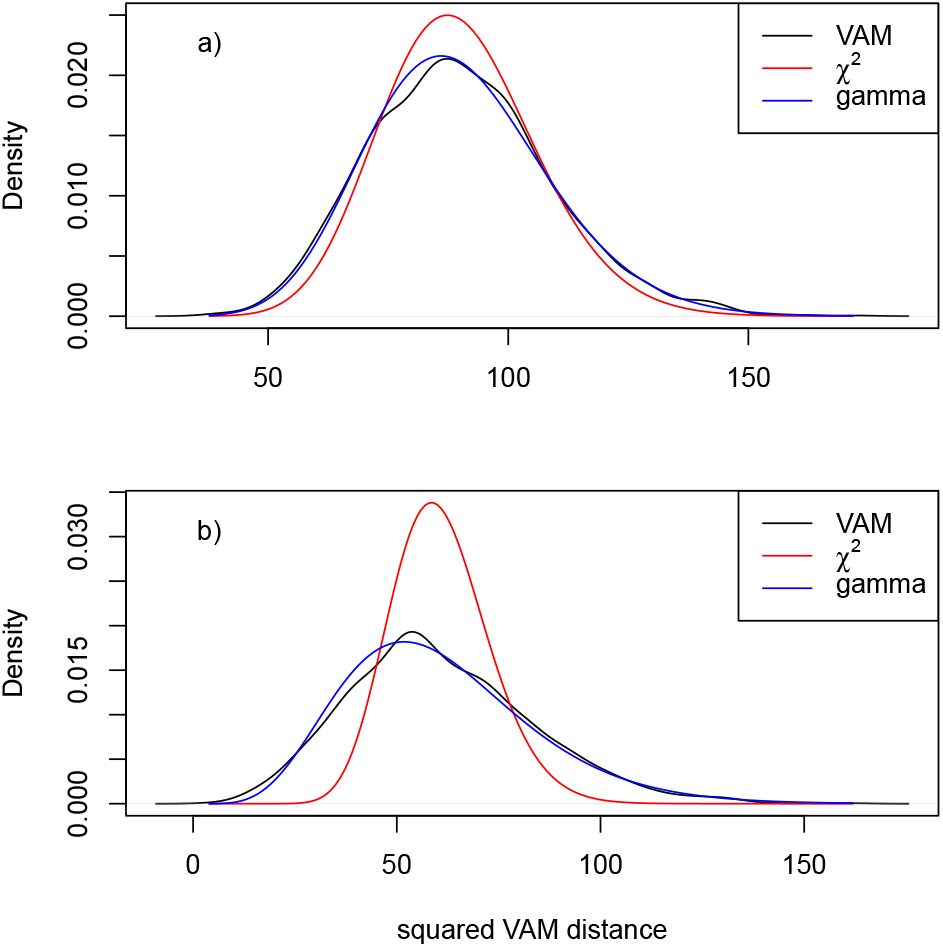
Distribution of squared modified Mahalanobis distances computed using (1) on scRNA-seq data simulated under the *H*_0_ of uncorrelated technical noise as detailed in Section 2.5. The densities of the non-central *χ*^2^ approximation and estimated gamma distribution are also plotted. **a)** Density estimates for data with a simulated sparsity of 0.5. **b)** Density estimates for data with a simulated sparsity of 0.8.

### 2.4 VAM-Seurat integration

The VAM implementation supports direct integration with the Seurat framework [35] with the integration details varying based on whether Seurat log-normalization is followed by variable feature detection using a mean/variance trend or the SCTransform [37] method is used to perform both normalization and variable feature detection. The **S** matrix generated by VAM is saved as a new Seurat assay, which enables the visualization and further analysis of these cell-specific pathways scores using Seurat framework, e.g., the *FeaturePlot()* and *FindMarkers()* functions.

#### 2.4.1 Integration for log-normalization

The Seurat log-normalization method implemented by the *NormalizeData()* R function starts by dividing the unique molecular identifier (UMI) counts for each gene in a specific cell by the sum of the UMI counts for all genes measured in the cell and multiplying this ratio by the scale factor 1 × 10^6^. The normalized scRNA-seq values are then generating by taking the natural log of this relative value plus 1. When log-normalization is used, variable features are detected using the *FindVariableFeatures()* function, which fits a non-linear trend to the log scale variance/mean relationship (the Seurat *vst* method). The estimated trend models the expected technical variance based on mean gene expression; observed variance values above this expected trend reflect biological variance. Given this trend, the proportion of technical variance is computed as ratio of the expected technical variance to the observed variance. Note that it is possible for this ratio to be greater than 1 if the observed variance is less that the expected variance. In this scenario, VAM sets the **X** matrix to the log-normalized values and the technical variance vector 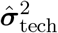 is computed as the product of the variance of the normalized counts and the proportion of technical variance as estimated by the *vst* method.

#### 2.4.2 Integration for SCTransform

The Seurat SCTransform normalization method [37] fits a regularized negative binomial regression model on the UMI counts for each gene using approximate cell sequencing depth as a dependent variable. The Pearson residuals from these regression models capture the biological component of the scRNA-seq data and should have a mean of 0 and variance of 1 if expression is due solely to technical noise. The reciprocal of the Pearson residual variance therefore estimates the proportion of technical variance. In this scenario, VAM sets the **X** matrix to 1 plus the natural log of the corrected UMI counts (i.e., counts that have been adjusted using the Pearson residuals to reflect the counts that would be observed if all cells had the same sequencing depth) and the technical variance vector 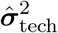 is computed as the ratio of the variance of corrected counts in **X** and the variance of the Pearson residuals.

### 2.5 Simulation study design

To explore the statistical properties of the VAM method, we used simulated scRNA-seq data simulated to reflect the characteristics of the PBMC log-normalized data. To simulate normalized scRNA-seq data, i.e, the contents of **X**, we took advantage of the fact that the non-zero log-normalized values in real scRNA-seq data sets can be effectively modeled by a log-normal distribution. Figure 3 illustrates this distributional approximation for the non-zero log-normalized counts from the peripheral blood mononuclear cell (PBMC) scRNA-seq data set (see Section 2.6 for more details on this data set). Based on this result, we simulated normalized scRNA-seq data under the *H*_0_ of un-correlated technical noise by first populating **X** for 2,000 cells and 500 genes with independent log-normal RVs with mean and variance set to the sample estimates for the non-zero normalized counts in the PBMC scRNA-seq data. The generated **X** was then sparsified by setting a random selection of elements to 0, with the number of zero elements matching the desired sparsity level. Data sets simulated according to this procedure were used to generate Figure 2 as well as the type I error control results in Section 3.1.

**Figure 3:**
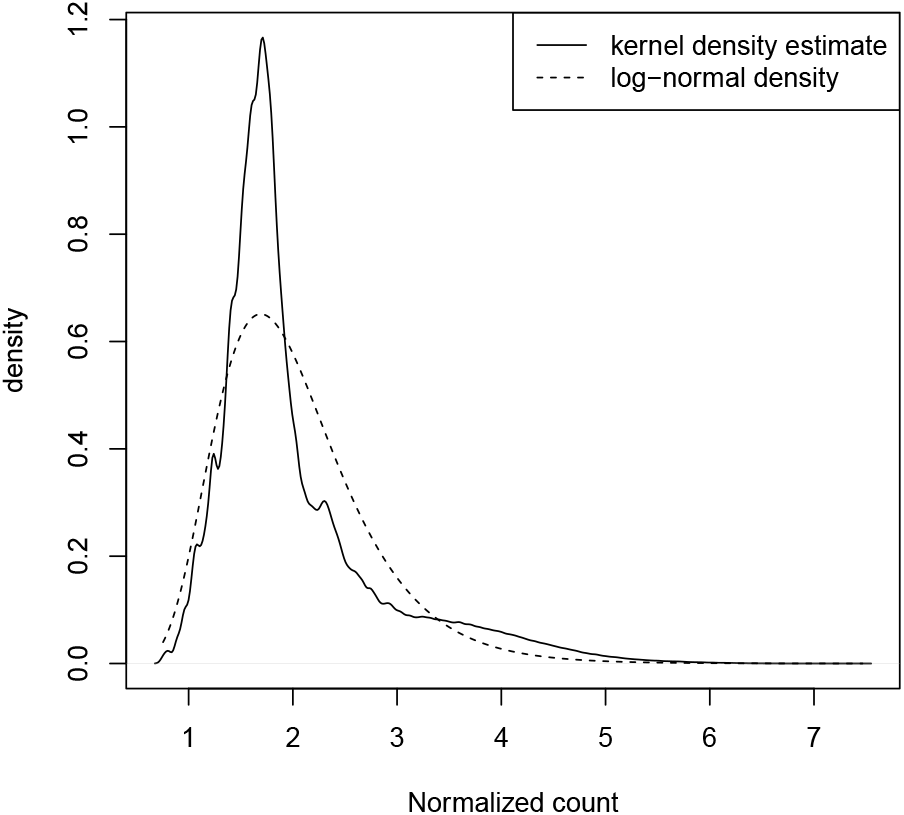
Distribution of non-zero log-normalized counts from the PBMC scRNA-seq data set. Both the kernel density estimate and associated log-normal density are displayed.

To assess power and classification performance (Sections 3.1 and 3.2), data sets simulated under *H*_0_ were modified to elevate the normalized expression of genes in a hypothetical gene set of size 50 for the first 50 cells while preserving the overall sparsity level. The elevated values were computed by setting all non-zero counts in the original null data to log-normal RVs with a larger mean (the variance was set to the same value used to simulate the null data). Classification performance was assessed for a range of null variance, set size and inflated mean values.

### 2.6 Real data analysis design

To assess the performance of VAM on real scRNA-seq data, we analyzed two data sets that are both freely available from 10× Genomics: the 2.7k human PBMC data set used in the Seurat Guided Clustering Tutorial [40], and an 11.8k mouse brain cell data set generated on the combined cortex, hippocampus and sub ventricular zone of an E18 mouse [41]. These two data sets are representative of small and medium sized single cell experiments and capture transcriptomic measurements for distinct heterogeneous cells populations (immune cells and neural cells) for the two organisms (human and mouse) that comprise a large fraction of existing scRNA-seq data sets. Preprocessing, quality control (QC), normalization and clustering of the PBMC data set followed the exact processing steps used in the Seurat Guided Clustering Tutorial. Specifically, the Seurat log-normalization method is used followed by application of the *vst* method for decomposing technical and biological variance. Preprocessing and QC of the PBMC data yielded an **X** matrix of normalized counts for 14,497 genes and 2,638 cells.

Processing of the mouse brain data followed similar quality control metrics (at least 200 features per cell, non-zero values in at least 10 cells for genes, proportion of mitochondrial reads less than 10% [42]) with Uniform Manifold Approximation and Projection (UMAP) [43] used for dimensionality reduction and clustering performed with Seurat’s implementation of shared nearest neighbor (SNN) modularity optimization [44]. Normalization of the mouse brain data was performed using SCTransform rather than log-normalization to explore the performance of VAM for both of the supported Seurat normalization approaches. Preprocessing and QC of the mouse brain data yielded an **X** matrix of normalized counts for 32,850 genes and 9,320 cells.

For the VAM analysis of these two scRNA-seq data sets, the gene set matrix **A** was populated using the C2.CP.BIOCARTA (BioCarta, 289 gene sets), and the C5.BP (Gene Ontology Biological Processes, 7,350 gene sets) collections from the version 7.0 of the Molecular Signatures Database (MSigDB) [38]. These MSigDB collections contain gene sets from three well known and widely used repositories of curated gene sets: BioCarta [38], and the biological process branch of the Gene Ontology [45]. Prior to running VAM, the Entrez gene IDs used by MSigDB were converted to Ensembl IDs using logic in the Bioconductor *org.Hs.eg.db* R package [46]. For analysis of the mouse brain data, the human Ensembl IDs were mapped to murine orthologs using logic in the *biomaRt* R package [47]. The **X** and **A** matrices were then filtered to only contain genes present in both matrices (13,714 genes for the PBMC data and 16,425 genes for the mouse brain data). Finally, the **A** matrix was filtered to remove all gene sets containing fewer than 5 or more than 200 members. To determine enrichment of gene sets for specific scRNA-seq clusters, a Wilcoxon rank sum test was performed using the Seurat *FindMarkers()* method.

### 2.7 Comparison methods

For comparative evaluation of the VAM method on both simulated and real scRNA-seq data, we used methods from each of the existing categories of single sample gene set testing methods. For the random walk category, we used both GSVA [27] and ssGSEA [28] given the popularity of these two techniques, for the class of z-scoring methods, we used the technique of Lee et al. [31], and, for the class of PCA-based methods we used PLAGE [30]. For all of these comparison methods, the implementations available in the GSVA R package were employed. Unless otherwise noted, analyses were performed using default values for method parameters.

## 3 Results

### 3.1 Type I error control and power

Type I error control was assessed using scRNA-seq data simulated according to the process detailed in Section 2.5 with the technical variances set to the sample variance of the simulated genes. The VAM method was applied to a set comprised by 50 randomly selected genes. The type I error rate at an *α* = 0.05 for 10 simulated scRNA-seq data sets (2,000 p-values per data set for 20,000 total hypothesis tests) was 0.048. To assess power, a random group of 50 genes were given inflated log-normal values for the first 50 cells with the mean value ranging from 0.7 to 1.7 (the non-inflated mean was 0.642 to align with the PBMC data). For each inflated mean value, 10 data sets were simulated and power was computed on the 50 non-null cells for a total of 500 hypothesis tests. As displayed in Figure 4, the estimated power values ranged from 0.11 for an inflated mean of 0.7 to 0.99 for an inflated mean of 1.7.

**Figure 4:**
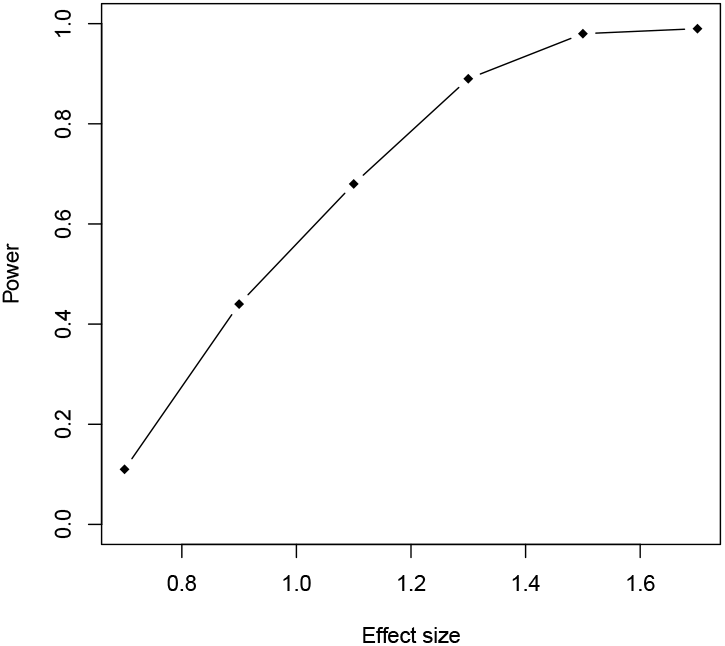
Estimated statistical power for the VAM method on simulated scRNA-seq data at different effect size values.

### 3.2 Classification performance

To compare the performance of VAM against existing single sample gene set testing methods, we measured the classification accuracy of each method (i.e., how well the method is able to highly rank cells that have inflated values for the genes in a specific set) on scRNA-seq data sets simulated according to the procedure outlined in Section 2.5. Use of classification accuracy vs. statistical power for the comparative evaluation had two motivations: 1) VAM is the only method in the comparison group that generates valid p-values, and 2) we envision VAM being used primarily as a means to rank order cells according to pathway activity rather than as a tool for cell-level statistical inference. Figure 5 illustrates the relative classification performance (as measured by the area under the receiver operating characteristic curve (AUC)) of VAM, GSVA [27], ssGSEA [28], and representative methods from the z-scoring and PCA-based categories (the technique of Lee et al. [31] for z-scoring and PLAGE [30] for PCA-based methods) across a range of sparsity, noise, effect size and set size values.

**Figure 5:**
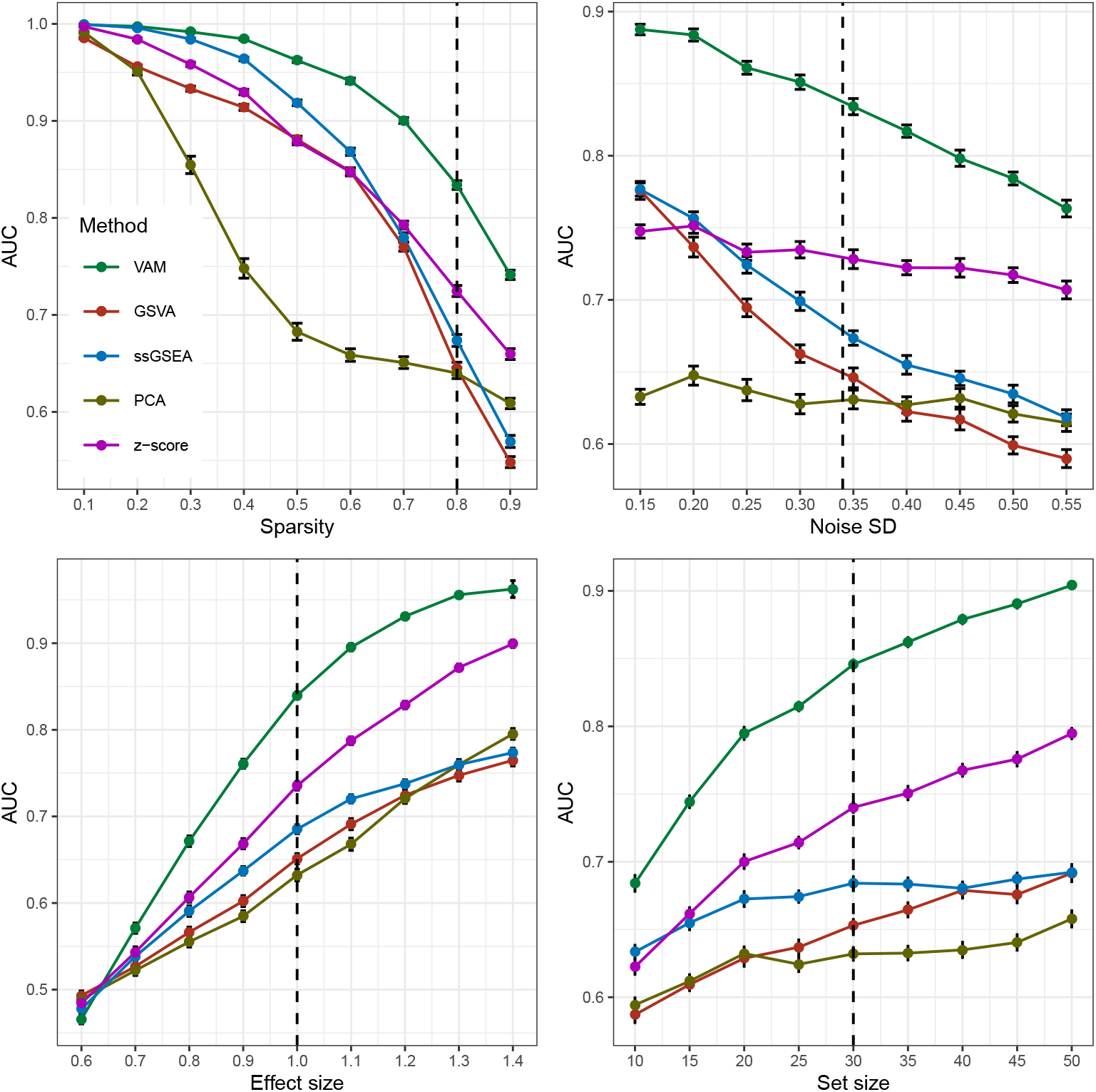
Classification performance of VAM, GSVA, ssGSEA and representative z-scoring and PCA-based methods on scRNA-seq data simulated according to Section 2.5. Each panel illustrates the relationship between the area under the receiver operating characteristic curve (AUC) and one of the simulation parameters. The vertical dotted lines mark the default parameter value used in the other panels. Error bars represent the standard error of the mean.

For each distinct combination of parameter values, 50 data sets were simulated according to the procedure outlined in Section 2.5 and Figure 5 displays the average AUC for each method across these 50 data sets with error bars representing the standard error of the mean. The general trends in performance follow the expected trajectories, e.g., AUC values fall as sparsity or noise increase and AUC values increase as the effect size or set size increases. Importantly, the VAM method provides superior classification performance relative to the other evaluated methods across the full range of evaluated parameter values with the difference particularly pronounced for the sparsity and variance found in the PBMC scRNA-seq data.

### 3.3 Computational efficiency

Table 1 displays the relative execution time GSVA, ssGSEA and representative z-scoring G and PCA-based methods as compared to VAM. Relative times are shown for the analysis of the simulated data sets (2,000 cells and 500 genes) used to generate the classification results shown in Figure 5, for the analysis of the PBMC scRNA-seq data set using the MSigDB BioCarta (C2.CP.BIOCARTA) collection (see Section 3.4 for detailed results on the PBMC data set), and for the analysis of the mouse brain scRNA-seq data set using the MSigDB Gene Ontology biological process (C5.BP) pathway collection (see Section 3.5 for detailed results on the mouse brain data set). A specific result for the GSVA method on the mouse brain data is not available since this method failed to complete the analysis due to memory issues. The VAM method had a much faster average execution on the simulated data set relative to the other methods with the difference particularly dramatic for the two most popular single sample methods, GSVA and ssGSEA. Although the PCA-based method was faster than VAM on the PBMC data and both the z-scoring and PCA-based methods were faster than VAM on the mouse brain data, the difference in execution time between VAM and both GSVA and ssGSEA on these real data sets was still over an order-of-magnitude.

**Table 1:** Relative execution time as compared to the VAM method on simulated scRNA-seq data, the PBMC scRNA-seq data set for MSigDB C2.CP.BIOCARTA collection and the mouse brain scRNA-seq data set for the MSigDB C5.BP collection. The GSVA method failed to process the mouse brain data so a specific relative performance is not available.

|  | Simulated | PBMC | Mouse brain |
| --- | --- | --- | --- |
| GSVA | 426.29 | 50.85 | - |
| ssGSEA | 23.99 | 22.68 | 26.61 |
| z-scoring | 6.14 | 3.06 | 0.23 |
| PCA | 2.63 | 0.38 | 0.62 |

**Table 2:**
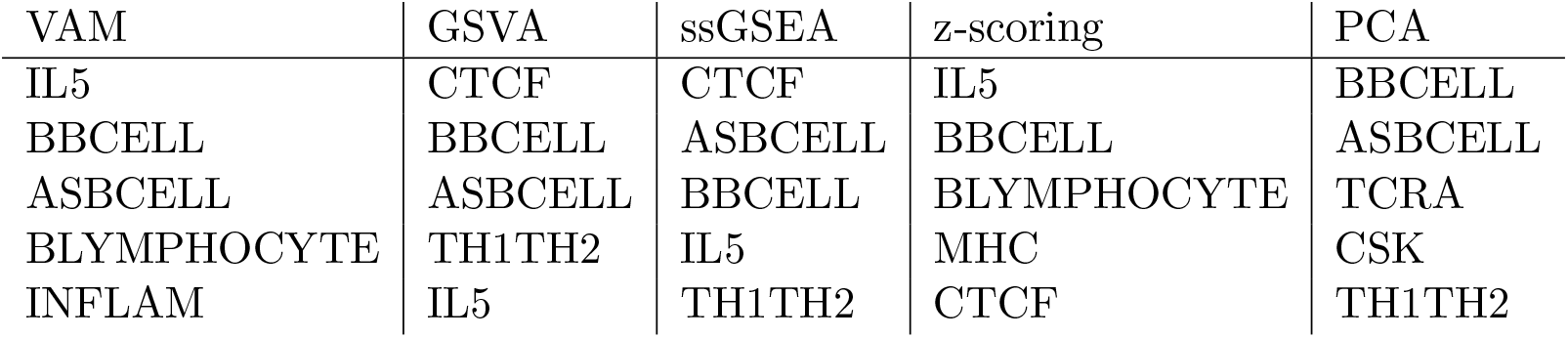
Top five BioCarta pathways found to have higher pathway activity scores in the B cell cluster relative to other cells in the PBMC data set according to a Wilcoxon rank sum test. Pathways are ordered according to p-value from Wilcoxon test. The columns reflect the method used to compute the cell-specific pathway scores.

### 3.4 Human PBMC analysis

As detailed in Section 2.6, we applied the VAM method and comparison techniques to the 10× 2.7k human PBMC data set used in the Seurat Guided Clustering Tutorial [40]. Figure 6 is a reduced dimensional visualization of the 2,638 cells remaining after quality control filtering. Cluster cell-type labels match the assignments in the Seurat Guided Clustering Tutorial. For this analysis, the cell-specific pathway scores were used to identify pathways with elevated activity within cell-type specific clusters. As an illustrative example, we highlight the results for the B cell cluster. Table 2 lists the five MSigDB BioCarta pathways most significantly up-regulated in the B cell cluster according to a Wilcoxon rank sum test applied to the cell-specific scores computed by VAM and other comparison methods. All of the evaluated methods correctly associate B cell-related pathways with the B cell cluster, which is not surprising given the very distinct transcriptomic profile of B cells. While all of the methods offer similar classification performance in this scenario, VAM still has the benefits of low computational cost and support for cell-level inference. For more complex cell populations, e.g., the mouse brain scRNA-seq data detailed in Section 3.5, VAM appears to offer superior classification performance relative to the other techniques.

**Figure 6:**
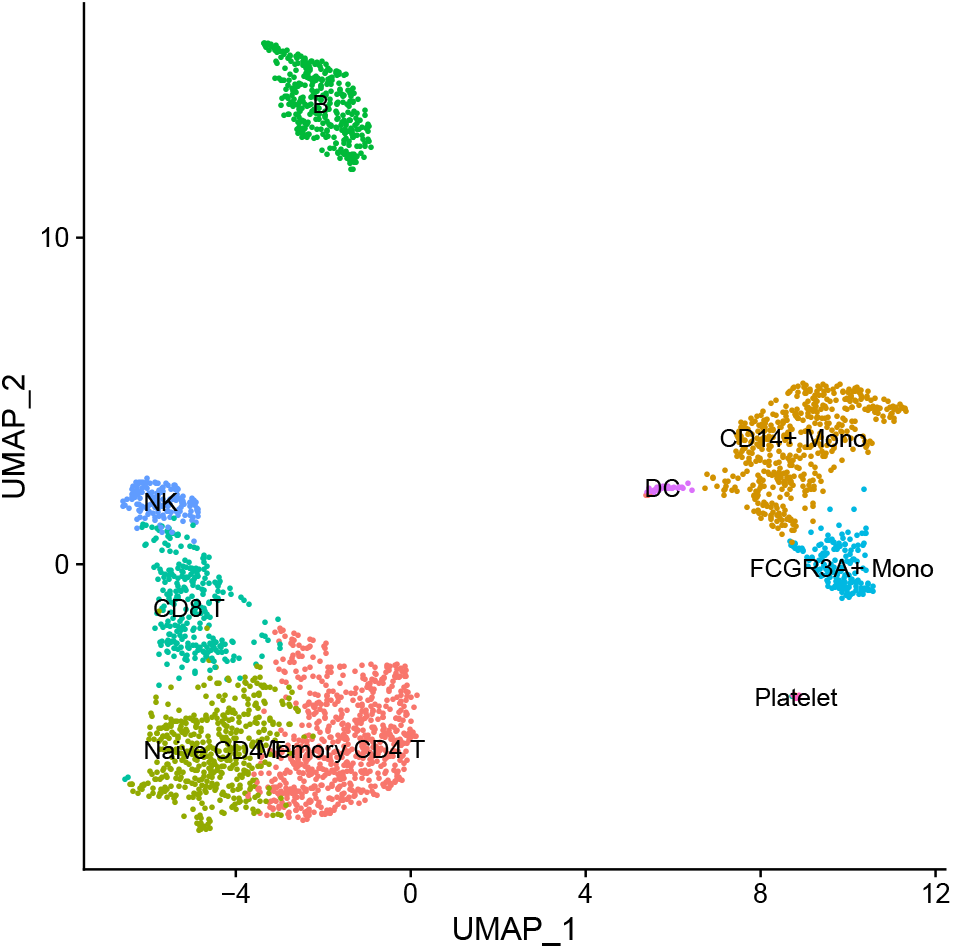
Projection of PBMC scRNA-seq data onto the first two UMAP dimensions. Each point in the plot represents one cell.

A important use for the cell-specific scores generated by VAM is the visualization of pathway activity across all cells profiled in a given scRNA-seq data set. Figure 7 illustrates such a visualization for the four BioCarta pathways most significantly up-regulated in the B cell cluster according to cell-specific scores generated by the VAM method. This type of visualization provides important information regarding the range of pathway activity across all profiled cells, e.g., IL-5 activity is also up-regulated in monocytes.

**Figure 7:**
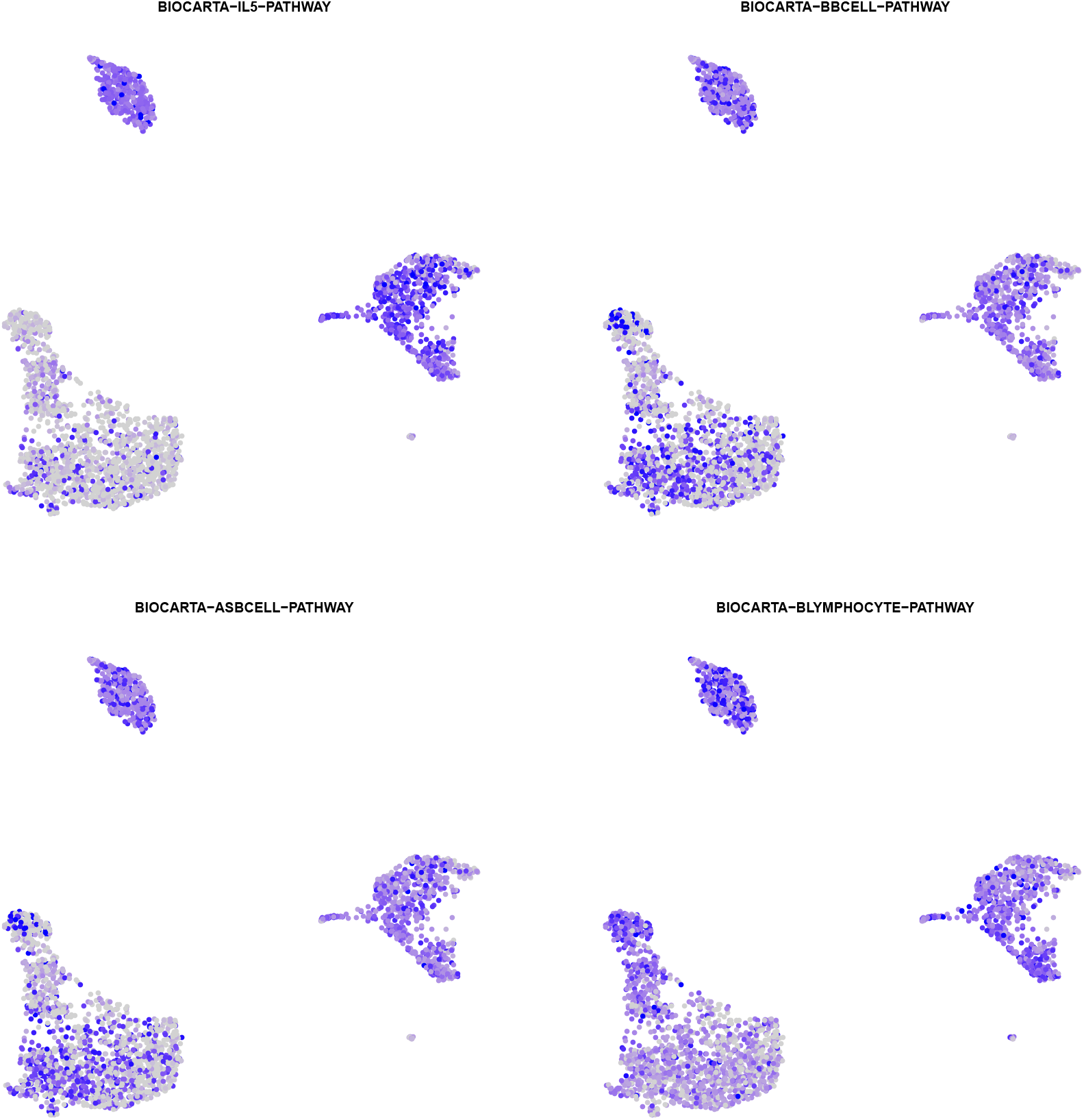
Visualization of the VAM generated cell-specific scores for the four BioCarta pathways most significantly enriched in the B cell cluster according to a Wilcoxon rank sum test on the VAM scores.

### 3.5 Mouse brain cell analysis

As detailed in Section 2.6, we applied the VAM method and comparison techniques to the 10× 11.8k mouse brain scRNA-seq data set. For this example, we used the SCTransform normalization technique instead of log-normalization and explored a much larger pathway collection (the MSigDB Gene Ontology biological process (C5.BP) collection with 7,350 gene sets). Figure 8 is a reduced dimensional visualization of the 9,320 cells remaining after quality control filtering with cells labeled according to the output from unsupervised clustering. Similar to the PBMC analysis, the cell-specific pathway scores were used to identify pathways with elevated activity within specific clusters.

**Figure 8:**
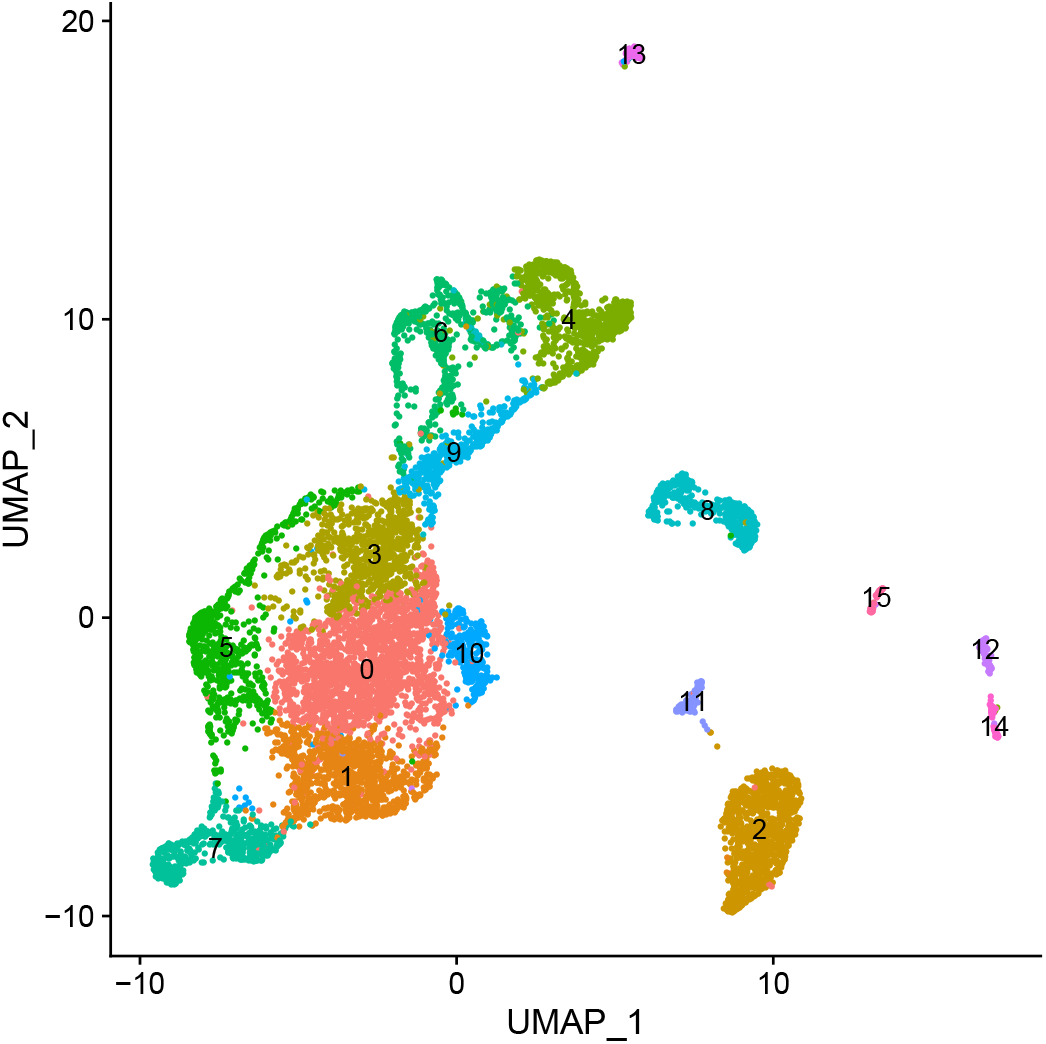
Projection of mouse brain scRNA-seq data onto the first two UMAP dimensions. Cells are labeled according to the output from unsupervised clustering.

We highlight the results for cluster 4, which appears to represent glial cells including a population of astrocytes, a glial cell subtype. Table 3 lists the five MSigDBC C5.BP gene sets most significantly up-regulated in cluster 4 according to a Wilcoxon rank sum test applied to the cell-specific scores computed by VAM and other comparison methods.

**Table 3:**
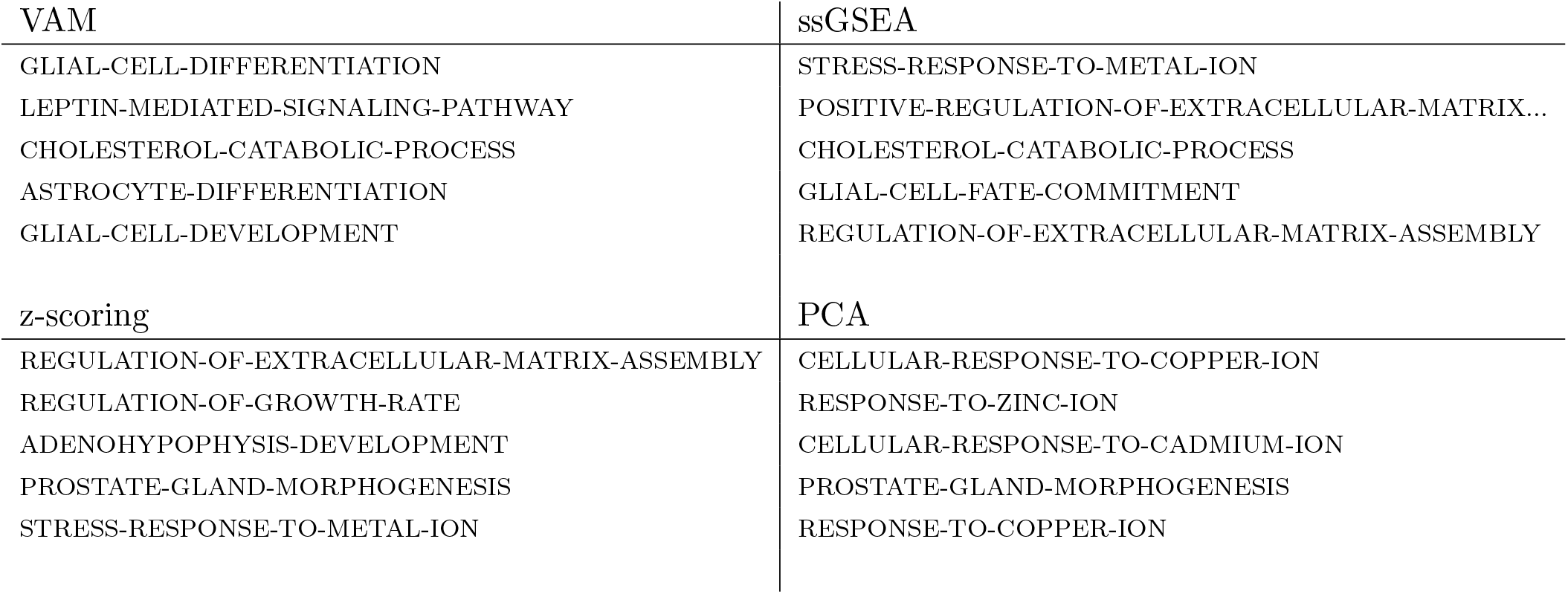
Top five Gene Ontology Biological Process gene sets (from MSigDB C5.BP collection) found to have higher pathway activity scores in cluster 4 relative to other cells in the mouse brain data set according to a Wilcoxon rank sum test. Gene sets are ordered according to p-value from Wilcoxon test. No results are available for the GSVA method since it failed to successfully process this data set.

As seen in Table 3, VAM clearly associates this cluster with glial cells with *GLIAL-CELL-DIFFERENTIATION* the top ranked set and both *ASTROCYTE-DIFFERENTIATION* and *GLIAL-CELL-DEVELOPMENT* also in the top five list. Figure 9 provides a visualization of the VAM-generated scores for the top four gene sets up-regulated in cluster 4. By contrast, neither the z-scoring nor PCA-based methods included glial cell-related sets in the top 5 and ssGSEA only identified one, *GLIAL-CELL-FATE-COMMITMENT*, at rank 4. None of these other methods identified an astrocyte-related gene set. Although it is not possible to say with certainty that cluster 4 captures the glial (and potentially astrocyte-specific) sub-population in this scRNA-seq data, the top five most significantly up-regulated genes in cluster 4 according to a Wilcoxon test on the SCTransform-corrected counts all have a known association with astrocytes: Dbi [48], Ptn [49], Tubb4b [50], Hopx [51], Igfbp2 [52].

**Figure 9:**
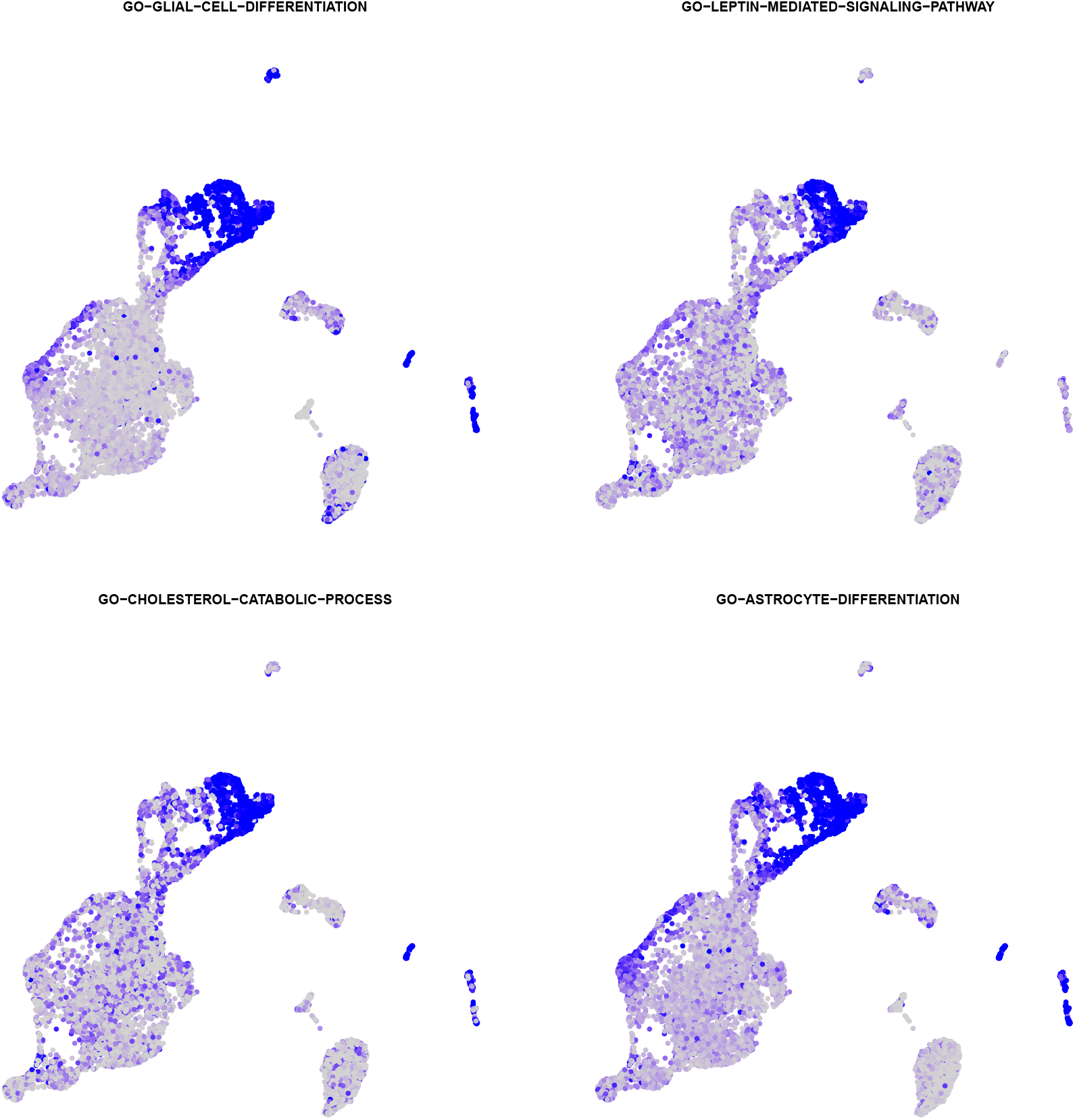
Visualization of the VAM generated cell-specific scores for the pathways most significantly enriched in cluster 4 (as seen in Figure 8) according to a Wilcoxon rank sum test on the VAM scores.

## 4 Conclusions

Single cell RNA-sequencing is a powerful experimental tool for exploring the biology of heterogeneous cell populations. The significant sparsity and technical noise associated with scRNA-seq data, however, makes statistical analysis challenging, especially for tests conducted on the level of individual genes. One promising approach for addressing the statistical challenges of scRNA-seq data is gene set testing or pathway analysis, a hypotheses aggregation technique that can mitigate the issues of sparsity and technical noise to improve power, replication and interpretability. The class of single sample gene set testing methods, which transform a cell-by-gene matrix into a cell-by-pathway matrix, is particular effective for single cell analyses since it enables the full range of standard downstream processing (visualization, clustering, differential expression testing, etc.) to be performed on the pathway-level rather than on the gene-level. Unfortunately, almost all existing single sample gene set testing methods were designed for the analysis of bulk tissue gene expression data, which is non-sparse and, compared to scRNA-seq data, has a small sample size and limited technical noise.

To remedy the lack of effective single sample gene set testing methods for scRNA-seq data, we developed the variance-adjusted Mahalanobis (VAM) method, a novel modification of the standard Mahalanobis multivariate distance measure that generates cell-specific pathways scores which account for the inflated noise and sparsity of scRNA-seq data. Although we expect the scores generated by VAM to be primarily used in contexts that do not assume a specific statistical model, e.g., as predictor variables, the fact that the distribution of the VAM-generated scores has an accurate gamma approximation under the null of uncorrelated technical noise enables inference regarding pathway activity for individual cells. As demonstrated on both simulated and real scRNA-seq data, the VAM method provides superior classification performance at low computational cost relative to existing single sample techniques. The utility of VAM is also aided by direct integration with the popular Seurat framework, which makes it easy to incorporate VAM into existing scRNA-seq analysis pipelines. These features combine to make the VAM method an effective and practical tool for the visualization and statistical analysis of scRNA-seq data.

